# A hybrid and poly-polish workflow for the complete and accurate assembly of phage genomes: a case study of ten przondoviruses

**DOI:** 10.1101/2023.03.09.531871

**Authors:** Claire K. A. Elek, Teagan L. Brown, Thanh Le Viet, Rhiannon Evans, David J. Baker, Andrea Telatin, Sumeet K. Tiwari, Haider Al-Khanaq, Gaëtan Thilliez, Robert A. Kingsley, Lindsay J. Hall, Mark A. Webber, Evelien M. Adriaenssens

## Abstract

Bacteriophages (phages) within the *Przondovirus* genus are T7-like podoviruses belonging to the *Studiervirinae* subfamily, within the *Autographiviridae* family and have a highly conserved genome organisation. The genome size of these phages ranges from 37 kb to 42 kb, encode 50-60 genes and are characterised by the presence of direct terminal repeats (DTRs) flanking the linear chromosome. These DTRs are often deleted during short-read-only and hybrid assemblies. Moreover, long-read-only assemblies are often littered with sequencing and/or assembly errors and require additional curation. Here, we present the isolation and characterisation of ten novel przondoviruses targeting *Klebsiella* spp. We describe HYPPA – a <u>HY</u>brid and <u>P</u>oly-polish <u>P</u>hage <u>A</u>ssembly workflow, which utilises long-read assemblies in combination with short-read sequencing to resolve phage DTRs and correcting errors, negating the need for laborious primer walking and Sanger sequencing validation. Our data demonstrate the importance of careful curation of phage assemblies before publication, and prior to using them for comparative genomics.

**IMPACT STATEMENT:** The current workflows employed for phage genome assembly are often error-prone and can lead to many incomplete phage genomes being deposited within databases. This can create challenges when performing comparative genomics, and may also lead to incorrect taxonomic assignment. To overcome these challenges we proposed HYPPA, a workflow that can produce complete and high-quality phage genomes without the need for laborious lab-based validation.

**DATA SUMMARY:** Phage raw reads are available from the National Centre for Biotechnology Information Sequence Read Archive (NCBI-SRA) under the BioProject number PRJNA914245. Phage annotated genomes have been deposited at GenBank under the accessions OQ579023-OQ579032 (**Table 1**). Bacterial WGS data for clinical preterm infant samples have been deposited at GenBank under BioProject accession PRJNA471164 (**Table S1**). Bacterial raw reads for food samples are available from NCBI-SRA with individual accessions (SAMN33593347-SAMN33593351), and can be found under the BioProject number PRJNA941224 (**Table S1**). Strain-specific details for bacteria and publicly-available phages used in these analyses, along with accessions for the latter can be found in **Table S1** and **Table S6**, respectively. The CL1-CL8 clinical *Klebsiella* strains (**Table S1**) were under a Materials Transfer Agreement, for which sequencing data and strain information is not available.

## INTRODUCTION

Double-stranded (ds) DNA bacteriophages with the characteristic head-tail morphology, also known as tailed phages, are a diverse group of viruses spanning 47 families, 98 subfamilies, and 1197 genera, with many more being unclassified (1-4). Phages within the *Przondovirus* genus are T7-like podoviruses, meaning they have a short tail morphotype, belonging to the *Studiervirinae* subfamily, within the *Autographiviridae* family (5). T7-like phages are renowned for following a strictly lytic life cycle, with the eponymous *Escherichia coli* phage T7 often used as the type isolate to represent the *Autographiviridae* family (6, 7).

*Autographiviridae* phages typically have genomes ranging from 37 to 42 kb in size and encode 50-60 genes, with the DNA-directed RNA polymerase (RNAP) being a hallmark of the family (5, 6, 8). The genome organisation of genera within the *Studiervirinae* subfamily is highly conserved: all genes are unidirectional and show a high degree of synteny (2, 5-7).

Tailed phages employ a remarkably diverse array of packaging methods that generate distinct termini (9, 10). The termini of T7-like phages consist of direct terminal repeats (DTRs) of varying lengths that flank the genome (6). The DNA of T7-like phages is concatemeric when generated within the bacterial cell and requires the assistance of terminases to cut at specific sites to package the DNA into the procapsid (9-11). Whilst each concatemer contains a single copy of the repeat, a second repeat is synthesised at the other end of the genome to prevent loss of genetic material (9, 12). Additionally, the DTRs are thought to prevent host-associated digestion *in vivo* and assist in DNA replication during phage infection (10, 13).

Many phage genomes deposited within public sequence databases are incomplete, often with DTR sequences missing or simply not annotated. Thus, our relatively limited understanding of phage biology is exacerbated by incomplete data and can make classification and comparative genomics more challenging (14). Indeed, high-quality genomic data will help identify relationships between taxonomic classification, infection kinetics, and phage-host interactions that are essential to the use of phages as therapeutics (14).

The genus *Klebsiella* comprises a heterogenous group of Gram-negative bacteria in the Enterobacterales order (15). *Klebsiella* spp. are common commensals of human mucosae, presenting a major risk factor for developing invasive disease and are therefore important opportunistic pathogens (15, 16). Antibiotic resistance among *Klebsiella* spp. represents a major threat to human health, with many isolates now multidrug resistant (15, 16). Therefore, conventional treatment using currently available antibiotics is becoming increasingly ineffective, and combined with no new antibiotics in the drug development pipeline, we are entering a post-antibiotic era (17, 18). Treatment of recalcitrant infections with bacterial viruses, bacteriophage therapy, has seen a resurgence in recent years as an alternative or adjunctive to current antibiotic therapy (19, 20).

Phage isolation involves monomicrobial or polymicrobial enrichment that often selects for the fittest phages (14, 21-23). Indeed, the rapid infection cycle of T7-like phages means that they are often overrepresented following traditional isolation methods (14, 21, 23). Here, ten novel T7-like phages belonging to the *Przondovirus* genus in the *Autographiviridae* family have been isolated against four *Klebsiella* strains belonging to different species, and characterised. Hybrid poly-polish assembly methods have recently been described for assembling bacterial genomes (24). Here we developed and validated a similar approach to ensure accurate and complete phage genome assembly, in a new worklow HYPPA – a <u>HY</u>brid and <u>P</u>oly-polish <u>P</u>hage <u>A</u>ssembly which was tested and validated for these new phages. The workflow utilises long-read assemblies in combination with short-read sequencing to resolve phage DTRs and correct sequencing and/or assembly errors, which negates the need for laborious primer walking and Sanger sequencing validation.

## MATERIALS AND METHODS

### Bacterial strains and growth conditions

Where specified, *Klebsiella* spp. used here were derived from previous studies (25-29) and are listed in **Table S1**. All *Klebsiella* strains were cultured overnight on brain heart infusion (BHI) agar (Oxoid) at 37°C. Liquid cultures were prepared by inoculation of 10 mL BHI broth with each bacterial strain and incubated at 37°C with shaking at 200 rpm for 3 h. Single colony variants were identified on solid media by changes in colony morphology and were purified by selecting a single colony for three successive rounds of purification on MacConkey no. 3 agar (Oxoid), and incubated overnight at 37°C.

### Preparation of bacterial DNA and sequencing

Genomic DNA for each *Klebsiella* strain was extracted using the AllPrep Bacterial DNA/RNA/Protein kit (QIAGEN) according to the manufacturer’s instructions. DNA was quantified as previously described and normalised to 5 ng µL^−1^.

DNA was prepared using the Illumina DNA Prep library preparation kit and was whole-genome sequenced on the Illumina NextSeq500 platform generating 2 × 150 bp paired-end reads by QIB Sequencing Core Services.

Additionally, *K. michiganensis* M7 21 2 #35, *K. pneumoniae* M26 18 1, *K. pneumoniae* M26 18 2 #21 KpnN, *K. pneumoniae* M26 18 2 #21 KpnA, and *K. pneumoniae* ST38 01 were prepared for enhanced sequencing according to the sequence facilities instructions (MicrobesNG). Genomes were provided assembled and annotated by MicrobesNG.

### Bacterial genomics

Short-read data provided without pre-processing by QIB Sequencing Core Services was QC filtered, trimmed, assembled, annotated, and analysed using the ASA^3^P v1.2.2 (30), or Bactopia v1.6.4 (31) pipelines. Preliminary strain designations were determined by ribosomal multilocus sequence typing (rMLST) (https://pubmlst.org/species-id) (32). The PubMLST database (https://pubmlst.org/) (33) was used to determine sequence type (ST) for *K. oxytoca* species complex and *K. aerogenes*, while the Institute Pasteur MLST database (https://bigsdb.pasteur.fr/) was used to determine sequence types of *K. pneumoniae* species complex. The capsular type for each strain was predicted using Kleborate (34) and Kaptive (35) on the QIB Galaxy platform, and those with a match confidence of good or higher were included.

### Isolation and single-plaque purification of phages

Samples from various UK wastewater treatment plants were screened for *Klebsiella*-specific phages using a range of *Klebsiella* strains as hosts for enrichment, adapted from Van Twest *et al*. (36). Briefly, 300 µL filtered wastewater was mixed with 60 μL exponential bacterial culture and used to inoculate 5 mL BHI broth. Enrichments were incubated overnight at 37°C with shaking at 200 rpm. Enrichments were then centrifuged (4000 x *g* for 15 min) and passed through a 0.45 μm filter before spot testing by double agar overlay plaque assay, as previously described (37). All incubations for overlay method were performed over 4-17 h at 37°C. Single plaque purifications were made by extracting single plaques from the soft agar layer using sterile toothpicks and suspended in approximately 300 μL BHI broth. Suspensions were centrifuged (13,000 x *g* for 5 min) and supernatant collected. Ten-fold serial dilutions of the supernatant were performed in phage buffer (75 mM NaCl; 10 mM MgSO_4_; 10 mM Tris, pH 7.5; 0.1 mM CaCl_2_) and 10 μL of each dilution plated onto double agar overlay and incubated as described above. This process was repeated at least three times to create phage stocks.

Phage amplification was performed as for single plaque purification in BHI broth. Once supernatant was collected, approximately 100 μL of phage suspension was spread onto three double agar overlay plates and incubated as before. Phage stocks were prepared by extraction of phage clearance zones. This was achieved by removal of the soft agar layer, which was resuspended in phage buffer, and centrifuged (4000 x *g* for 15 min). Phage supernatant was passed through a 0.45 µm filter into a sterile glass universal and stored at 4°C.

### Phage host range

Phage host range was tested by plaque assay as described above on a range of clinical, wastewater, food, and type strain *Klebsiella* spp. as described previously (38). Only assays where individual plaques were identified were recorded as positive.

### Phage DNA extraction and whole-genome sequencing

Phage virions were concentrated by polyethylene glycol (PEG) 8000 (Thermo Fisher) precipitation for DNA extraction. Briefly, phage stock was treated with 1 µL DNase I (10 U µL^−1^) (Merck) and 1 μL RNase A (10 U µL^−1^) (Merck) per mL of stock and incubated at 37°C for 30 min. PEG precipitation was performed with PEG 8000 (10% w/v) and 1 M NaCl and incubated overnight at 4°C. The precipitate was centrifuged (17,000 x *g* for 10 min) and resuspended in 200 µL nuclease-free water. Resuspended phage pellets were treated with proteinase K (50 µg mL^−1^) (Merck), EDTA (final concentration 20 mM), and 10% SDS (final concentration 0.5% v/v) and incubated at 55°C for 1 h.

DNA was extracted using the Maxwell® RSC Viral Total Nucleic Acid Purification kit (Promega), as per the manufacturer’s instructions into nuclease-free water. Phage DNA was quantified by Qubit 3.0 fluorometer using the high sensitivity dsDNA kit (Invitrogen). DNA was prepared using Illumina DNA Prep (formerly Nextera Flex) library preparation kit and was whole-genome sequenced on the Illumina NextSeq500 platform generating 2 × 150 bp paired-end reads by QIB Sequencing Core Services. MinION libraries (Oxford Nanopore Technologies, ONT) were constructed without shearing using the short fragment buffer and loaded onto the R9.4.1 flow cell according to the manufacturer’s instructions by QIB Sequencing Core Services.

Both long-read and short-read raw data for all ten przondoviruses were deposited in NCBI under BioProject number PRJNA914245.

### Phage genomics

#### Assembly and annotation

All quality control, pre-processing, assembly, and annotation of phage genomes were performed on the QIB Galaxy platform.

We checked short-read data for quality using fastQC v0.11.8 (39). Based on this fastQC analysis, reads were pre-processed with fastp v0.19.5 (40), using a hard trim of between 4 and 10 bases on both the front and tail to retain at least a per base quality of 28.

Long-read data was demultiplexed following sequencing and quality checked with NanoStat v0.1.0 (41). Pre-processing was performed as part of the assembly, and assembled using Flye v2.9 (42) with default settings, which included correction and trimming of reads. Flye was used in the first instance as previously published work has determined it is the most accurate and reliable assembler (43-45). Where Flye was unable to generate a high-quality assembly, Canu v2.2 (46) was used as an alternative. Error correction and trimming were performed as part of the default settings when assembling using Flye or Canu. Flye additionally performed one iteration of long-read polishing by default. We assembled all phages with and without trimming adapter/barcode sequences for long-reads. Trimming was performed with Porechop v0.2.3 (https://github.com/rrwick/Porechop) (47) with default settings.

We performed several iterations of long-read and short-read polishing on long-read-only assemblies in a specific order. Firstly, two iterations of long-read polishing were performed using Medaka (48) with default settings, using the previous polished data as the input for the next round of polishing. Secondly, one iteration of short-read polishing was performed using Polypolish (49) with default settings. Finally, a second iteration of short-read polishing was performed using POLCA (50) with default settings. We used raw reads for each iteration of long-read polishing and pre-processed reads for each iteration of short-read polishing.

Prior to development of the current phage assembly workflow, we had adopted a few other methodologies for resolving the genomes. One method was short-read-only assembly, where phages were assembled *de novo* using Shovill v1.0.4 (https://github.com/tseemann/shovill) with default settings (51, 52). Briefly, trimming was disabled by default and manual trimming was performed as part of the pre-processing step prior to assembly. Additionally, SPAdes was used as the default assembler within the Shovill pipeline. We attempted short-read polishing of long-read-only data using Pilon v1.20.1 (53) with default settings. Where specified, we also performed hybrid assembly using raw long-read and pre-processed short-read data, as previously described using Unicycler v0.4.8.0 (54) with default settings. Porechop v0.2.3 (https://github.com/rrwick/Porechop) (47) was used for *Klebsiella* phage Oda only. All assembly details are given in **Tables S3-S5**.

Following assembly, the contigs were manually checked for DTRs flanking the genome, as well as with PhageTerm (55) which was unable to identify the DTRs since it does not work well for Nextera-based sequence libraries. Where we could not determine the length and sequence of the DTRs, we performed primer walking. Outward-facing primers were designed to “walk” the genome termini using Sanger sequencing (56). Phage DNA was extracted, and for each phage at least two primers were designed for the reverse strand to walk the beginning of the genome and identify the left terminal repeat, and at least two primers were designed for the forward strand to walk the end of the genome to identify the right terminal repeat. The phage DNA and each primer were then sent for Sanger sequencing separately (Eurofins, Germany). Sanger sequences were visualised in FinchTV v1.5.0 (https://digitalworldbiology.com/FinchTV) and compared to the reference phage genome, and DTRs annotated using the Molecular Biology suite on the Benchling platform (https://www.benchling.com/).

Assemblies generating multiple contigs were checked for contamination using Kraken 2 v2.1.1 (57).

Verification of the DTRs and assessment of assembly quality was performed by mapping the raw reads back to the assembled genome using Bowtie 2 v2.3.4.3 (58) and visualised using IGV v2.7.2 (59), and variant calling performed using iVar v1.0.1 (60). Additionally, BWA-MEM v0.7.17.1 (https://github.com/lh3/bwa) was used to map long-reads back to the reference using default settings optimised for ONT reads (61, 62).

Assemblies in the reverse orientation were reorientated by reverse complementation of the genome in UGENE v38.0 (63) and uploaded to Benchling. Contigs were then reoriented to begin at the same start point, based on well-curated reference phages and the analysis of the DTRs.

Genome annotation was performed using Pharokka v1.2.1 with default settings (https://github.com/gbouras13/pharokka) (64). Specifically, coding sequences were predicted with PHANOTATE (65).

#### Comparative genomics

Where specified, publicly-available phage genomes used for comparative genomics were derived from these studies (20, 66-78), listed in **Table S6**, and downloaded from the GenBank database.

The closest relative for each phage was determined as as the top hit according to maximum score identified by nucleotide BLAST (BLASTn) (https://blast.ncbi.nlm.nih.gov/Blast.cgi) and optimised for somewhat similar sequences (79). Genes associated with specific phage families were identified and used for preliminary taxonomic assignment. Alignments were performed using Mauve v20150226 (80) between the closest relative and phages from the same genera. The intergenomic similarity between przondoviruses in the collection and a selection of publicly-available related phages was calculated using VIRIDIC on the web server (http://rhea.icbm.uni-oldenburg.de/VIRIDIC/) (81).

Phylogenetic analyses were performed using the hallmark DNA-directed RNA polymerase (RNAP) amino acid sequence for all phages and a selection of publicly-available phylogenetically-related phages downloaded from the NCBI protein database (https://www.ncbi.nlm.nih.gov/). Multisequence alignment of the RNAP amino acid sequences was performed using the MUSCLE algorithm in MEGA X v10.0.5 (82) with default settings. A maximum-likelihood tree was generated with 500 boostraps using the default Jones-Taylor-Thornton model. Phylogenetic analysis was performed using 35 amino acid sequences, with a total of 684 positions in the final analysis. Tree image rendering was performed using iTOL v6.1.1 (https://itol.embl.de/) (83).

Linear mapping of coding sequences for phage final assemblies was performed using Clinker v.0.0.23 (84).

## RESULTS AND DISCUSSION

### Phage isolation and host range determination

In this study, we isolated ten lytic T7-like phages from a variety of river water and wastewater samples, using four different *Klebsiella* spp. as isolation hosts (**Table 1**). To examine the host range, we tested the ten phages against a collection of *Klebsiella* spp. from different sources, representing a range of capsule and sequence types. All phages had a narrow host range, with seven being able to infect only a single *Klebsiella* strain within our collection (**Fig. 1**).

**Table 1.** Przondoviruses within the collection to date and data relating to the closest database relative.

| Phage Name | Source | Isolation host | Genome size (bp) | GC content (%) | DTR size (bp) | No. CDS | Accession | Closest database relative according to BLASTn |  |  |
| --- | --- | --- | --- | --- | --- | --- | --- | --- | --- | --- |
|  |  |  |  |  |  |  |  | Name | Coverage (%) | ID (%) |
| Oda | River water | K.mi. | 41,642 | 52.64 | 181 | 58 | OQ579023 | *Klebsiella* phage SH-KP152226 | 92.0 | 94.62 |
| Toyotomi | Wastewater | K.mi. | 41,268 | 52.64 | 180 | 55 | OQ579024 | *Klebsiella* phage SH-KP152226 | 92.0 | 94.71 |
| Mera | Wastewater | K.mi. | 41,400 | 52.58 | 180 | 56 | OQ579025 | *Klebsiella* phage SH-KP152226 | 92.0 | 94.34 |
| Speegle | Wastewater | K.mi. | 41,395 | 52.64 | 180 | 58 | OQ579026 | *Klebsiella* phage SH-KP152226 | 93.0 | 94.70 |
| Cornelius | Wastewater | K.mi. | 40,437 | 52.72 | 180 | 55 | OQ579027 | *Klebsiella* phage SH-KP152226 | 94.0 | 94.84 |
| Tokugawa | Wastewater | K.mi. | 41,414 | 52.64 | 181 | 56 | OQ579028 | *Klebsiella* phage SH-KP152226 | 92.0 | 94.70 |
| Saitama | Wastewater | K.qp. | 40,741 | 53.06 | 181 | 51 | OQ579029 | *Klebsiella* phage K11 | 96.0 | 95.71 |
| Emom | Wastewater | K.ox. | 40,788 | 52.56 | 183 | 53 | OQ579030 | *Klebsiella* phage KP32 | 94.0 | 93.14 |
| Amrap | Wastewater | K.ox. | 41,209 | 52.47 | 182 | 57 | OQ579031 | *Klebsiella* phage KPN3 | 85.0 | 95.05 |
| Whistle | Wastewater | K.va. | 40,735 | 52.40 | 181 | 54 | OQ579032 | *Klebsiella* phage IME264 | 94.0 | 94.78 |

**Fig. 1.**
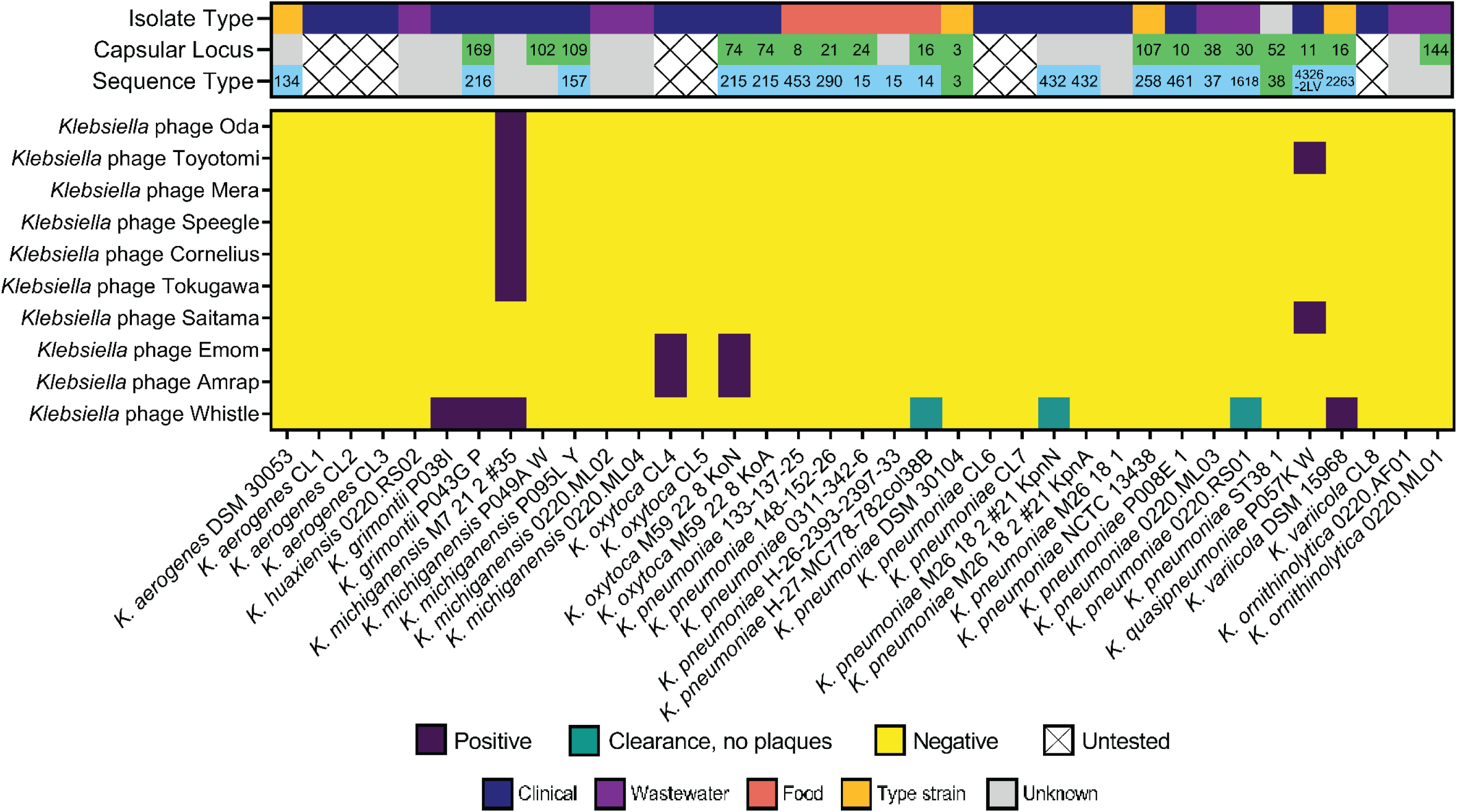
Heatmap for host range of the przondoviruses in the collection by plaque assay against a diverse range of *Klebsiella* spp. Top panel, isolate type, capsular type, and sequence type. The source of each isolate is given as isolate type, with grey being unknown source. Capsular loci determined by Kaptive and/or Kleborate, green; unknown or no match confidence, grey. Sequence type (ST) determined by multilocus sequence typing, blue; unknown or incomplete matches, grey. No sequencing data available, untested. Bottom panel, host range heatmap. Productive infection (positive) is the observation of individual plaques, purple; lysis without productive infection is the observation of clearance without individual plaques, green; no productive infection or clearance (negative), yellow.

Three of the ten przondoviruses were used to test and validate the HYPPA workflow: Oda, Toyotomi, and Tokugawa. As the three unifiers of the HYPPA workflow, these were named after the three unifiers of Japan (see **Development of a new workflow for the assembly of complete phage genomes**).

Only three of our phages were capable of productively infecting more than one *Klebsiella* strain. *Klebsiella* phage Toyotomi was able to infect two different species; a *K. michiganensis* strain (its isolation host) and a *K. quasipneumoniae* strain. *Klebsiella* phages Emom and Amrap were both able to infect two different isolates of *K. oxytoca. Klebsiella* phage Whistle was the only phage capable of productively infecting four strains of *Klebsiella*, spanning three different species (*K. grimontii, K. michiganensis*, and *K. variicola*), and caused lysis without productive infection on a further three *K. pneumoniae* isolates. We could not establish a link between capsular type and host range for these phages.

Przondoviruses and other T7-like phages have a relatively small genome of 37 to 42 kb, and this may limit their host expansion capabilities (for taxonomic assignment of the ten phages in this study, see section **Phage genome characterisation and taxonomy**). However, Whistle was capable of infecting multiple hosts, along with Emom, Amrap, and Toyotomi. Previous work has shown that T7-like phages are capable of infecting multiple hosts (66) and that host range is determined by interaction between phage receptor binding proteins, i.e. tail fibre and/or spike proteins, and bacterial cell receptors (14, 66, 85). LPS components are almost always identified as the secondary receptor for irreversible attachment in Gram-negative-targeting podoviruses (6, 14). Whether initial interaction with the outer membrane and degradation of the CPS constitutes a *bone fide* reversible attachment step, or whether this is a prerequisite to reversible attachment by the phage to another outer membrane component is yet to be fully elucidated (6, 14, 86, 87).

Some phages can be “trained” to increase their host range through co-evolution assays (19, 88). This may be particularly useful in cases of lysis from without, such as observed in Whistle, as they are already capable of binding to host receptors but unable to cause productive infection.

Intriguingly, Toyotomi was the only phage capable of infecting two different *Klebsiella* species, that none of its closest relatives from our collection were capable of infecting, despite exceptionally high protein sequence similarity across their tail fibre proteins.

Multiple factors affect host range and broadly involve extracellular and intracellular mechanisms. Extracellular mechanisms involve the ability of phages to bind to specific phage receptors on the bacterial cell surface that facilitate DNA ejection (89). Intracellular mechanisms involve evasion of phage defence systems that facilitate phage propagation (89). Expression of diffusible depolymerases facilitate interaction of phages with their primary and secondary receptor. This extracellular mechanism is more likely to explain the ability of Whistle to infect more than one isolate since there is productive infection. Thus, the ability several przondoviruses in our collection to infect different *Klebsiella* isolates could indicate that they share similarities in the chemical composition of their capsules, enabling degradation by a single depolymerase and allowing access to the phage receptors on the bacterial cell. Moreover, the bacterial isolates could share similar sugar motifs within their LPS structures, which are thought to be the secondary receptor of phages within the *Autographiviridae* family (6).

### Development of a new workflow for the assembly of complete phage genomes

To generate complete and accurate genomes for these ten phages, which included resolving the defined ends of phage genomes, and correcting sequencing and/or assembly errors, we utilised a long-read-only assembly with sequential polishing steps. This methodology exploited both long-read and short-read sequencing data in a workflow that we have named HYPPA – <u>HY</u>brid and <u>P</u>oly-polish <u>P</u>hage <u>A</u>ssembly (see also **Materials and Methods**) before moving onto annotation and comparative genomics (**Fig. 2**). Firstly, the long-reads were assembled using Flye or Canu, followed by two iterations of long-read polishing with Medaka. Next, we performed two iterations of short-read polishing using Polypolish (for the first iteration) and POLCA (for the second iteration).

**Fig. 2.**
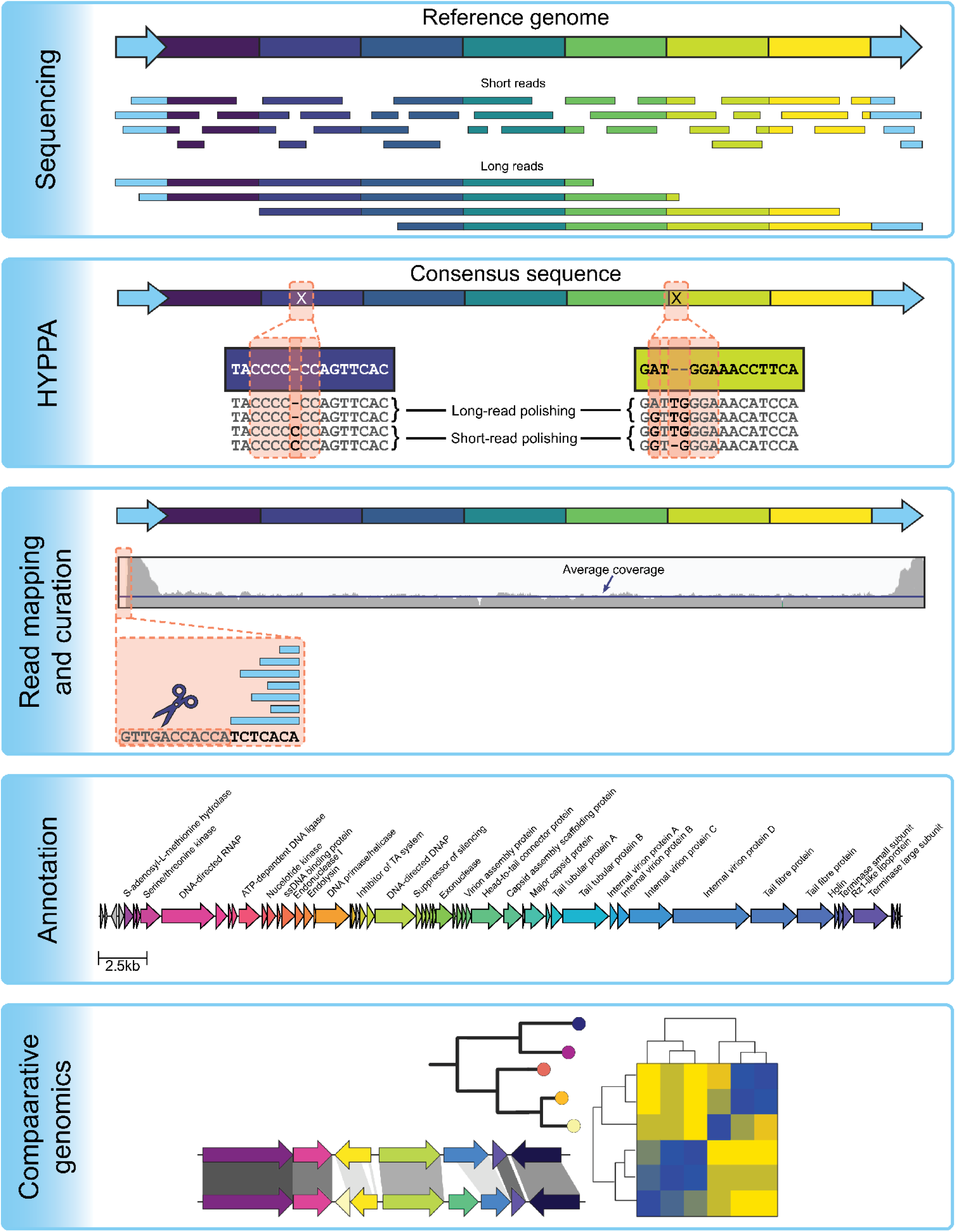
Workflow for phage genome assembly and analysis. Both long-read and short-read sequencing are recommended, followed by our HYPPA workflow for high quality phage genome assembly. Read mapping for short-read data and manual curation can correct any errors that were missed during polishing, followed by annotation and then comparative genomics.

Initially, Flye was used as the primary assembler in our HYPPA workflow and worked particularly well for phages with both very high sequence read coverage (Toyotomi at >117,000x) and very low sequence read coverage, which included Mera (8x), Speegle (23x), and Amrap (27x) (**Table S2**). However, Canu performed better with the other phages as the assemblies in general contained fewer errors. This is contrary to previously published literature that found Flye was more the more accurate assembler using default settings (43-45).

As an illustration of the HYPPA workflow, we provided a more detailed description of the process for phage Oda as an exemplar, for which the DTRs were validated with primer walking. Firstly, Oda was assembled using Canu, which yielded one contig of 41,761 bp. After two iterations of long-read polishing followed by two iterations of short-read polishing, the resulting contig was 41,769 bp in size. We were able to identify the terminal repeat regions, but both were flanked by a 64 bp sequence upstream of the left terminal repeat, and downstream of the right terminal repeat after all polishing iterations were complete. The two 64 bp sequences were inverted repeats containing adapter sequences of 23 bp, with the remaining sequence being Nanopore barcodes which were manually removed. HYPPA was then used for phage Tokugawa, which after short-read only assembly had included a 79 bp repeat within the genome, but outside of the presumed DTR region (**Fig. S1**). Using HYPPA, the repeat was determined to be an assembly artefact and removed from the assembly. The final curated assembly for phages Oda and Tokugawa was 41,642 bp and 41,414 bp, respectively. Terminal repeats were present for both phages and complete at 181 bp, validated by primer walking and Sanger sequencing (**Fig. S1**).

We trimmed the long-reads using Porechop in an attempt to remove the adapter/barcode sequences, but when phage Oda was reassembled and polished using the trimmed reads, the right terminal repeat was missing three bases, but no other single nucleotide polymorphisms (SNPs) or indels were identified.

The HYPPA workflow without Porechop-mediated trimming was repeated for the remaining eight przondoviruses, resulting in final genome assemblies ranging between 40-42 kb (**Table 1**). HYPPA was able to generate a complete genome for phage Toyotomi, where short-read-only, long-read-only, and hybrid assemblies were unable to do so and resulted in fragmented assemblies. Although our HYPPA workflow is a hybrid assembly approach, there is a clear distinction between this and traditional hybrid assembly methods. Importantly, HYPPA used the short-reads for polishing only, not during the genome assembly, whereas traditional hybrid assemblies utilise both long-read and short-read data during the assembly process itself. Moreover, short-read polishing of a long-read-only assembly using Pilon was also unable to resolve the genome of Toyotomi: partial repeat regions were found at the termini but were incomplete, and multiple errors within coding regions persisted. Using HYPPA, we were able to not only preserve the DTRs of Toyotomi, but also correct persistent sequencing and/or assembly errors that occurred in all non-HYPPA assemblies.

The genome organisation of genera within the *Autographiviridae* family is highly conserved: all genes are unidirectional and show a high degree of synteny, and genomes are flanked by DTRs (2, 5-7). The DTRs of the przondoviruses described here were 180-183 bp in size, demonstrating sequence similarity of 84.3-99.7%. DTRs are thought to assist circularisation of the phage genome once in the host cytoplasm to prevent host-induced enzymatic digestion (13). Thus, resolution of the DTRs is integral to accurate genomics and understanding of the biology of different phages.

### Comparison of HYPPA with traditional short-read-only assembly

When compared to typical short-read-only methodologies of phage genome assembly, in our case using Shovill (51), the HYPPA workflow required significantly less manual curation (**Fig. 2**). Typically, phage genomes are assembled using short-read only data, and many of these genomes are then published without additional curation, leaving them with potentially significant sequencing and/or assembly errors. Using short-read-only assembly methods for our collection of przondoviruses, we observed that some were in the reverse orientation rather than the forward orientation as is expected for 50% of the assemblies, and some had the DTRs assembled in the middle of the contig. Addressing these issues required manually re-orienting the assemblies and ensuring they all had the same start position, as suggested in the Phage Annotation Guide (90). In contrast, the HYPPA workflow resulted in assemblies with correct start and stop sites, but some were still in the reverse orientation.

To check for DTRs in short-read only assemblies, we initially looked for increased reads within the read mapping profiles, which are distinguished by one or two large peaks, and can be automated using the tool PhageTerm (55). If a single peak was observed anywhere other than at either end of the assembly, the assembly had been opened in the middle of the genome and required each to be re-oriented to have the same starting position.

Incorrect orientation is a feature of phage genome assembly, and with short-read-only data in particular, may be artificially linearised by the assembler with the DTRs located in the middle of the contig. In many of our own short-read-only assemblies, the przondoviruses described here were linearised in the middle of the genome, and required read mapping to identify where the DTRs may be. In T7-like phages, DNA is concatemeric and requires the assistance of terminases to cut at specific sites to package the DNA into the procapsid (9-11). Although each concatemer contains a single copy of the repeat, a second repeat is synthesised at the other end of the genome to prevent loss of genetic material (9, 12). Since the DTRs are present twice per phage genome, the number of terminal sequences is double following whole genome sequencing and are identified as a single peak of increased reads during read mapping (10-12, 55). Therefore, the DTR and by proxy, the start of the genome can be inferred from the read mapping. Moreover, due to the highly conserved nature of the genomes, all przondoviruses had almost the same starting sequence as the well-curated Enterobacteria phage K30 (accession HM480846) (67), making the beginning relatively easy to find. As a result, considerable time was spent on re-orienting the short-read-only assemblies to be unidirectional and to have the same starting sequence.

One of the most problematic aspects using short-reads for phage assembly (both short-read-only and has part of a traditional hybrid assembly) was that the DTRs were deleted, possibly because the assemblers used deem them to be a sequencing artefact. Thus, DTRs need to be manually validated through primer walking and Sanger sequencing validation. However, this was unnecessary when using short-reads for polishing rather than for assembly. Thus, using the HYPPA workflow, the DTRs were present in the final polished assembly in the correct location at the ends and did not have to be manually added.

A second type of error that routinely occurred during non-HYPPA phage sequencing and assembly was the introduction of short insertions and/or deletions (indels) that were particularly noticeable in coding regions.

For the short-read only assemblies, many sequencing and assembly errors present in coding regions were only found upon annotation of the genomes, including frameshift errors in DNA polymerase (DNAP) and tail fibre protein genes. Often, these frameshift errors were found in homopolymer regions and were introduced during sequencing. Before using HYPPA, these frameshift errors were checked through read mapping followed by variant calling and edited accordingly. Particularly noteworthy were repeat regions of ∼79 bp identified close to and sometimes within the DTR regions of seven of the ten phages (See **Development of a new workflow for the assembly of complete phage genomes** for description of repeats for Tokugawa), but that did not correlate with the increased reads observed in the read mapping. This suggested that these repeats were introduced in error during assembly and were confirmed to be artefacts in most phages, including Tokugawa through Sanger sequencing (see supplementary **Fig. S1**). Using HYPPA, we found that the two iterations of short-read polishing were able to correct single nucleotide polymorphisms and/or correct indels that resulted in these frameshift errors that long-read polishing was unable to resolve, particularly in homopolymer regions. POLCA was also able to correct indels that Polypolish was unable to resolve.

As previously described for Oda, all the przondoviruses contained adapter and barcode DNA upstream and/or downstream of the DTR regions. Initially, as we were trying to reconstruct the linear genome ends, we did not perform adapter and barcode trimming of the Nanopore reads prior to the long-read assembly. We then removed these sequences manually after assembly. To limit the amount of manual curation, Porechop can be used to trim the reads, however, when we attempted this for all the remaining przondoviruses, Porechop-mediated trimming resulted in several further errors. These included trimming bases from the beginning of the left terminal repeat and the end of the right terminal repeat, ranging from 3-18 bp in total; indels; multiple SNPs; and in some cases failure to assemble the phage genome into a single contig, or at all. We would thus recommend manual removal of the adapter/barcodes rather than trimming of long-reads using Porechop, which appears to require more manual curation when compared to using raw Nanopore reads.

Multiple sequencing and/or assembly errors were identified in the coding regions of other phages, that again, persisted following traditional methods of phage assembly. Using trial and error, we were able to show that the HYPPA method was superior to other methods of phage assembly, whether hybrid or through using a single sequencing platform in correcting errors (see supplemental **Tables S2-S5** for all assembly details). Moreover, the HYPPA workflow required far fewer manual curation steps than traditional phage assembly methods: while long-read only assemblies were sometimes in the reverse orientation, all were linearised at the starting sequence. This is in contrast with the traditional assembly methods that required re-orienting the genomes to be unidirectional and starting at the same position, manual correction of large assembly errors such as indels, manual correction of homopolymer errors in coding regions, and in some cases, rearrangement of contigs and manual stitching the genome together, followed by primer walking and Sanger sequencing validation to determine the genome termini and DTRs.

Errors in homopolymer sequences and repeat regions are particularly common in long-read-only assemblies of bacterial genomes (43, 44, 49), and as we have described here, in phage genomes also. Indeed, two homopolymer errors occurred in the DNAP of Toyotomi, leading to a double frameshift error that resulted in three protein annotations. Short-read polishing can correct errors introduced during long-read-only assemblies (49), as we have demonstrated here. Similarly to using short-read data for polishing, we found that a traditional hybrid assembly using both short-and long-read data for Toyotomi also introduced large deletions in repeat regions, with assembly errors persisting, as has been described previously (44, 54). Assembly metadata showing all previous long-read-only, short-read-only, and hybrid assemblies is provided (see supplemental **Tables S2-S5**).

Several limitations of this study include the need for both short-read and long-read data for phage assembly, and specialised knowledge to access and install the software which is all freely available. Which polishing program used and what type of polishing (long-read versus short-read) in what order may give different results of equal validity. While we believe that the HYPPA workflow provides the most accurate phage genome possible, it still may not exactly reflect the DNA that is present within each phage capsid. Additionally, while the highly conserved nature of T7-like phages made it easier to determine the DTR starting sequence, this may not be the case for novel phages.

### Phage genome characterisation and taxonomy

All ten phages were dsDNA phages at 40,336-41,720 bp with a GC content of 52.40-53.06% which is slightly lower than their isolation host GC content of ∼55.46-57.59% (**Table 1, Fig. 3**). The number of predicted coding sequences within the genomes varies from 51 to 58, and almost all coding sequences were found in the same orientation on the forward strand. However, five phages had one to four small hypothetical proteins found in opposite orientation. We performed BLASTn on all phages to determine closest relatives in the NCBI GenBank database (as of October 2022). Based on the BLASTn results, which showed high levels of nucleotide similarity with reference phages, the phages in our collection were preliminarily assigned to the *Przondovirus* genus within the *Studiervirinae* subfamily and *Autographiviridae* family, according to the currently established ICTV genus demarcation criterion of 70% nucleotide sequence similarity over the genome length to belong to the same genus (1).

**Fig. 3.**
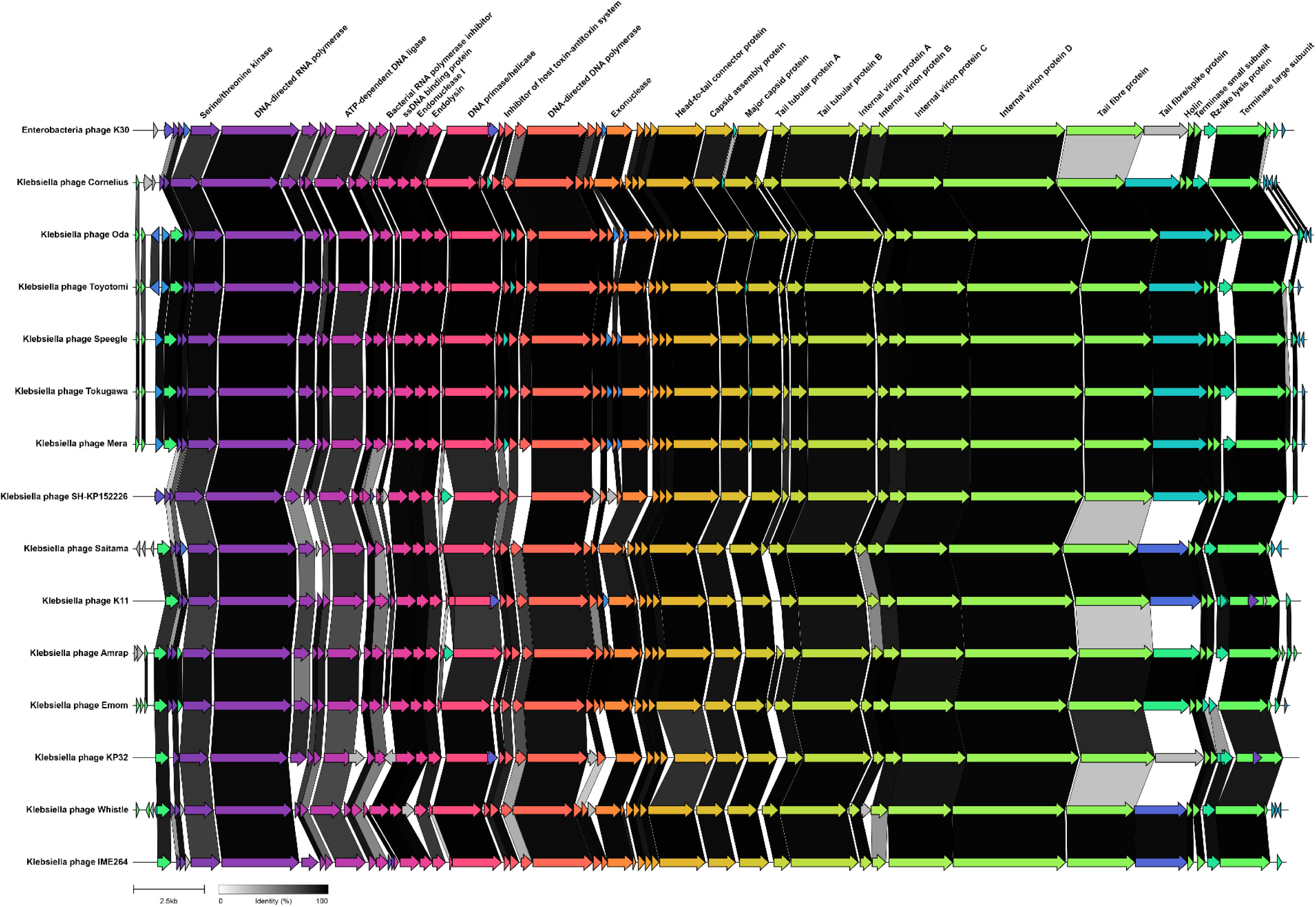
Genome map and gene clustering for przondoviruses in the collection and a selection of related phages. Arrows represent coding sequences and pairwise comparisons of gene similarities are indicated by percentage identity given as links in greyscale, with darker shading representing areas of higher similarity. Genes without any sequence similarity are indicated without links. Some phages had a hypothetical protein following the tail fibre protein and protein BLAST revealed high homology to tail spike proteins. DTRs are present but not annotated.

The genomic relationships between our novel przondoviruses and a selection of *Autographiviridae* reference phages were explored further by conducting a nucleotide-based intergenomic similarity analysis using VIRIDIC (**Fig. 4, Table S2**). Included within the analysis were relatives within the same genus (*Przondovirus*), those within different genera but the same subfamily (*Studiervirinae*), and those within different subfamilies (*Molineuxvirinae, Slopekvirinae*) (**Fig. 4**). These data confirmed that the przondoviruses from this study were within the ICTV genus demarcation criterion of 70% nucleotide sequence similarity over the genome length when compared to other przondoviruses. Several genera within the subfamily *Studiervirinae* that were included only shared ∼45-57% nucleotide sequence similarity with the przondoviruses in this study (**Fig. 4**).

**Fig. 4.**
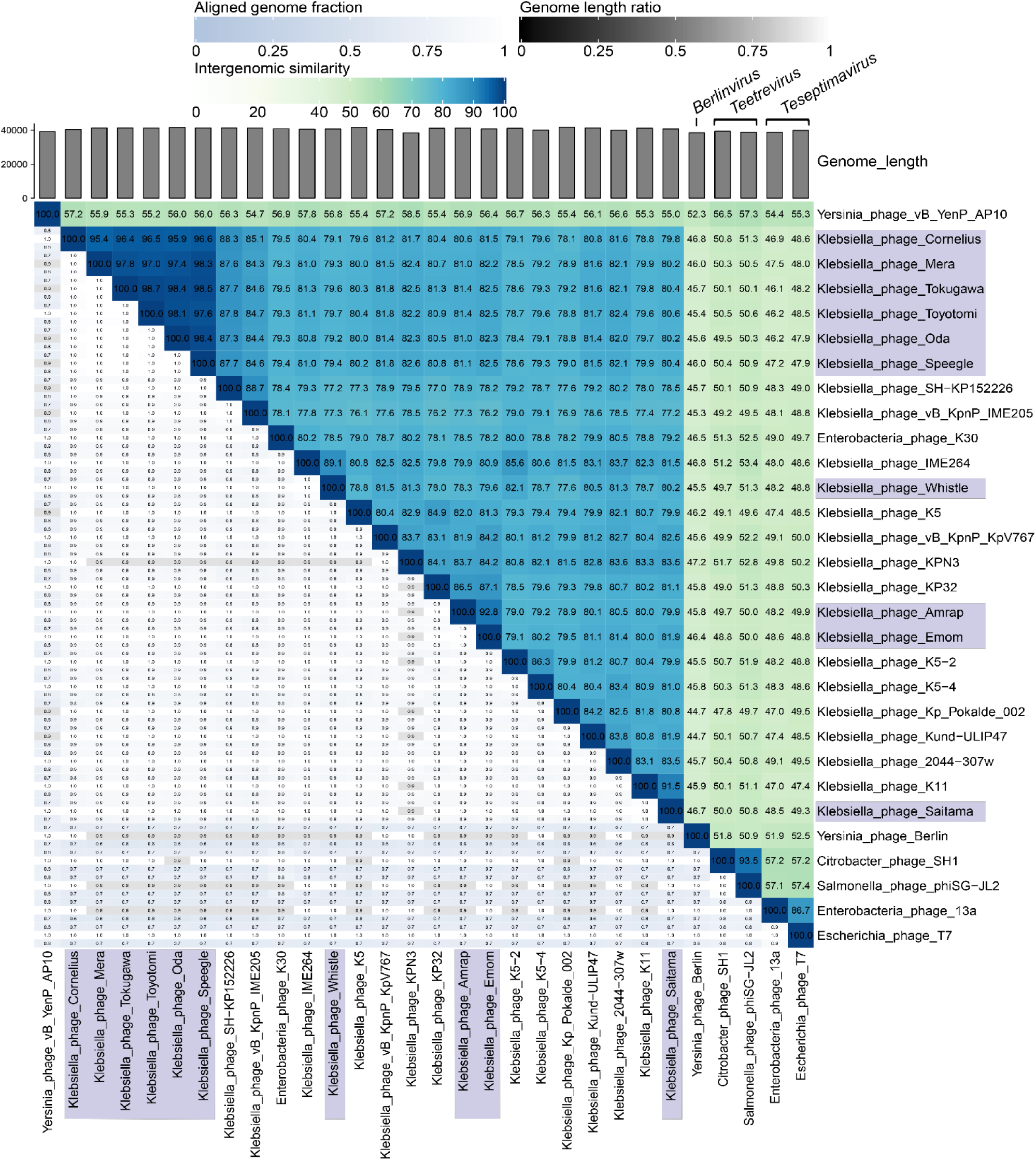
Nucleotide-based intergenomic similarities of przondoviruses in the collection and a selection of related phages within the *Studiervirinae* subfamily, using VIRIDIC. A heatmap of hierarchical clustering of the intergenomic similarity values was generated and given as percentage values (right half, blue-green heatmap). Each genome pair is represented by three values (left half), where the top and bottom (blue scale) represent the aligned genome fraction for the genome in the row and column, respectively, where darker colour indicates a lower fraction of the genome was aligned. The middle value (grey scale) represents the genome length ratio for each genome pair, where darker colour indicates increasing distance between phages. The przondoviruses within our collection are highlighted in blue-grey. Yersinia phage vB_YenP_AP10 is in the *Apdecimavirus* genus.

Several przondoviruses clustered more closely together, including *Klebsiella* phages Oda, Toyotomi, Mera, Speegle, Cornelius, and Tokugawa, which were within ∼98% nucleotide similarity, except Cornelius which was the most dissimilar at ∼95-96% (**Fig. 4**). All aforementioned phages except Oda were isolated from the same wastewater treatment plant at different stages of the treatment process, using the same host. These phages are therefore likely to be different strains of the same new species of phage within the *Przondovirus* genus. Emom and Amrap clustered with their closest relative KP32, but also clustered together with ∼92% similarity, and should be assigned to separate species (**Fig. 4**). Saitama and Whistle did not cluster closely with any other phage from our collection, possibly due to differences in their host specificity. Saitama did cluster with its closest relative *Klebsiella* phage K11, and Whistle clustered with its closest relative IME264 (**Fig. 4**). This suggests that Saitama, Emom, Amrap, and Whistle should be assigned to different species within the same genus.

After comparative genomic analyses, we observed that several of the closest database relatives were deposited in databases with incomplete genomes. Specifically, the incompleteness was most often due to an absence of the DTRs, including *Klebsiella* phages KP32, KPN3, and IME264 (**Table S2, Fig. S2**.). Incomplete genomes could lead to incorrect assignments to species in cases where the reciprocal nucleotide identities are close to the species threshold of 95% similarity across the genome length (1).

Additionally, potential errors were noted in phages KPN3 (accession MN101227) and KMI1 (accession MN052874) (**Table S2, Fig. S2**). For example, KPN3 contained no annotated DNA-directed DNAP, which is conserved across all *Przondovirus* genomes analysed here. KMI1 contained a shorter DNA-directed RNAP annotation that, when included in the phylogenetic analyses, showed higher divergence, which could not be confirmed, and was therefore excluded from our phylogenetic analysis. Without raw short-read and long-read data, it is difficult to determine whether these are genuine errors or whether their differences are a true representation of the genome.

To further verify the taxonomic classification of the phages, phylogenetic analysis was performed using the protein sequence of the DNA-dependent RNAP, since it is the hallmark gene of the *Autographiviridae* family, using a selection of publicly available phages from the genera *Apdecimavirus, Berlinvirus, Przondovirus, Teetrevirus*, and *Teseptimavirus*, within the subfamily *Studiervirinae* (**Fig. 5**). As expected, the pzondoviruses clustered together, and there was a clear separation from other phage genera. There were some slight differences between the clustering patterns exhibited in the phylogenetic tree when compared to the VIRIDIC analysis using whole nucleotide data. *Klebsiella* phages Emom and Amrap exhibited relatively high similarity, sharing 91.5% sequence similarity across the whole nucleotide sequence, but this distinction is less obvious in the phylogenetic analysis. As observed previously (91), a single gene phylogenetic tree at the amino acid level will not provide enough resolution to accurately display within-genus relationships, leading to discrepancies between clustering of the phages Saitama, Emom, Amrap and Whistle in the VIRIDIC plot and the phylogenetic tree.

**Fig. 5.**
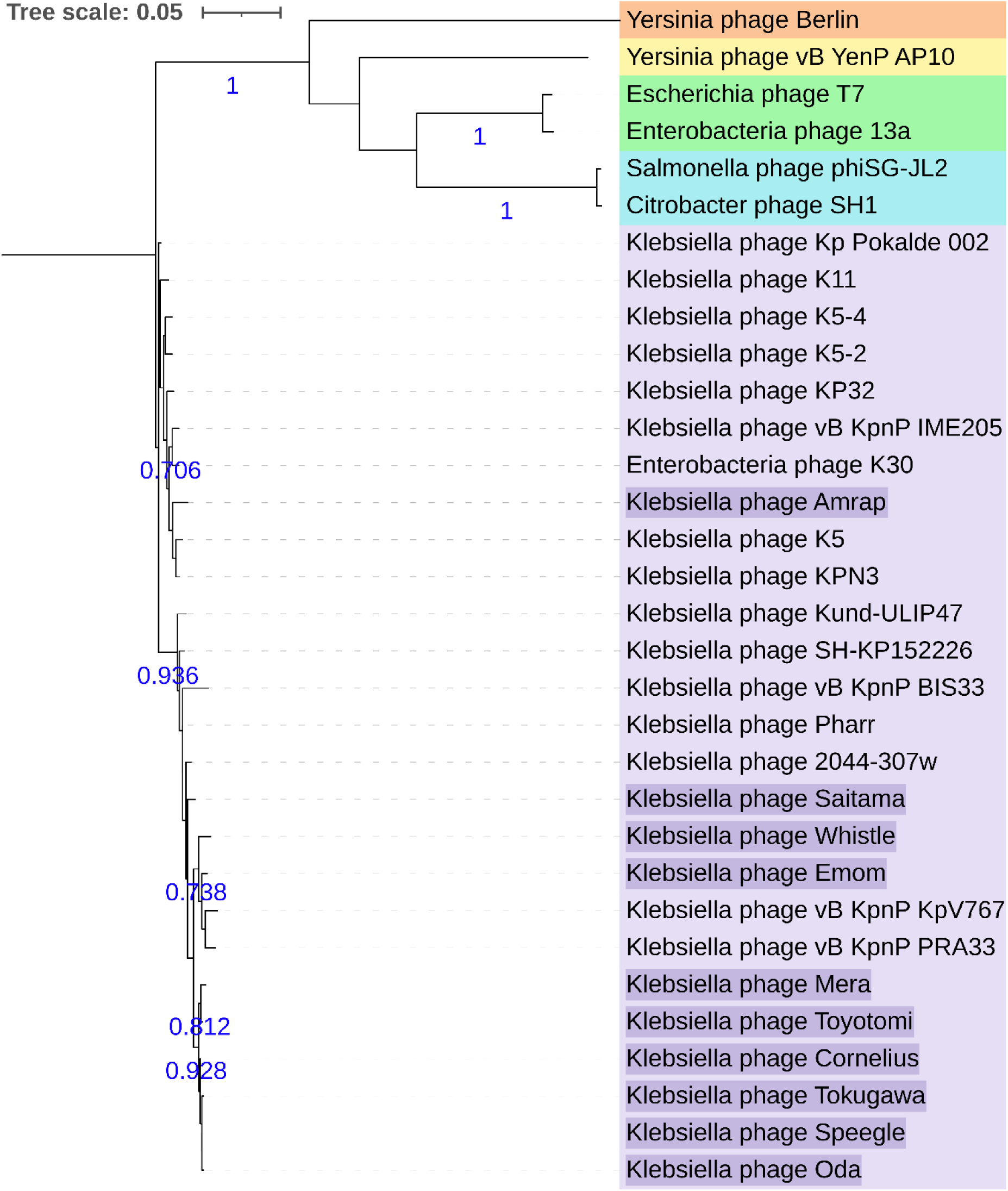
Maximum-likelihood phylogeny of the RNAP for przondoviruses in this study and a selection of related *Studiervirinae* phages. All phages from this study (purple highlight) clustered with related przondoviruses (purple). Outgroups, *Berlinvirus* (orange), *Apidecimavirus* (yellow), *Teseptimavirus* (green), and *Teetrevirus* (blue). Tree is midpoint rooted. Bootstrap support values at ≥0.7 are given in blue (500 replicates).

### Genome organisation and synteny

We conducted comparative genomic analysis of przondoviruses according to coding sequence similarity with a selection of reference phages (**Fig. 3**). We selected Enterobacteria phage K30 as the representative isolate of the *Przondovirus* genus since its genome is well-curated. Przondoviruses were grouped together with their closest relative according to BLASTn. As expected, all phages share a highly conserved genome organisation, which revealed a high degree of gene synteny, in concordance with the VIRIDIC data (**Fig. 4**).

All genomes were found to contain the early, middle, and late genes associated with viral host takeover, DNA replication, and virion assembly and lysis, respectively (**Fig. 3**). The host takeover proteins that were annotated included the S-adenosyl-L-methionine hydrolase, which is a good marker for the start of the genome; serine/threonine kinase; and DNA-directed RNAP, with the latter being a hallmark of the *Autographiviridae* family (5, 8). The middle proteins annotated were typical for phage DNA replication. The late proteins included all the components necessary for virion assembly, such as capsid proteins and tail-associated proteins, and lysis such as holins and Rz-like lysis proteins. Of the tail-associated proteins, two tail fibre and/or spike proteins were annotated for each przondovirus.

Within the *Przondovirus* genus, the main differences were found in the tail proteins (**Fig. 3**). The tail fibre and tail spike proteins are major determinants for host range, so phages that were isolated against the same *Klebsiella* host strain were expected to have higher sequence similarity across their tail fibre proteins. *Klebsiella* phages Oda, Toyotomi, Mera, Speegle, Cornelius, and Tokugawa, which were isolated against the same *K. michiganensis* strain, shared considerable sequence similarity across their entire genomes, including the tail fibre proteins. However, one difference was that Toyotomi was able to infect two hosts, whereas the remaining phages were not. This is intriguing since the high homology in the tail fibre protein would suggest similar narrow host range capabilities for this subset of przondoviruses in our collection. Emom and Amrap were both isolated against the same *K. oxytoca* strain, where they shared sequence similarity across their entire genomes, including at the tail fibre protein location. The tail fibre protein sequence similarity is complemented by the host range data for these two phages. In contrast, Cornelius and its closest relative *Klebsiella* phage SH-KP152226 still shared a high degree of sequence similarity across their entire genome, including the tail proteins, despite infecting different host species (*K. michiganensis* and *K. pneumoniae*, respectively). In fact, all przondoviruses in this study were found to share significant sequence similarity in their tail proteins with their closest relatives, except for Emom, and by proxy Amrap, and *Klebsiella* phage KP32. There was a lower degree of sequence similarity in the first tail fibre protein between Emom and KP32, but there was no sequence similarity in the second tail protein between Emom and that of KP32. This is possibly due to their different isolation hosts, where KP32 had been isolated against a *K. pneumoniae* strain, and Emom/Amrap were isolated against a *K. oxytoca* strain.

The most striking differences however, were in the tail proteins between przondoviruses in this study and reference phages that were not their closest BLASTn relatives. For example, Saitama showed sequence similarity with SH-KP152226 in only the initial part of the first tail fibre protein, with no sequence similarity exhibited elsewhere in the tail protein location. A similar pattern was observed for Emom and K11, and Whistle and KP32. This is unsurprising since the isolation host for Emom and K11 are *K. oxytoca* and *K. pneumoniae*, respectively (69, 92). Similarly, Whistle and KP32 infected two different species, *K. variicola* and *K. pneumoniae*, respectively. The differences in the tail fibre proteins therefore likely reflect the different isolation hosts for the przondoviruses in our collection and their database relatives.

Other differences between the closely related phages were found in the Rz-like lysis proteins, particularly within the przondoviruses that were within 95-98% similarity to one another. There is high sequence similarity for this protein between Cornelius and Oda, but not between Oda and Toyotomi, for example. Rz-like lysis proteins are involved in the lysis of the inner and outer membrane of Gram-negative bacteria and can be highly diverse (93-95). These proteins may be part of a single-component system, or part of a two-component system: this is where one gene may be embedded within another, overlap another, or exist as separate genes (93-95). These genes encode two different proteins that operate together to disrupt the bacterial membrane, but appear to have distinct evolutionary origins (95). The differences in membrane composition among different *Klebsiella* spp. could explain the differences in the Rz-like proteins, or may simply highlight differences between not only the proteins themselves, but the type of lysis system employed by each phage.

## CONCLUSION

Here, we developed the HYPPA workflow for generating high quality phage genomes that require minimal manual curation, and is most representative of what is actually biologically present within the phage capsid. We tested and validated the workflow using ten przondoviruses, negating the need for laborious primer walking and Sanger sequencing validation. Accurate phage genomes provide the necessary foundation for a mechanistic understanding of infection biology, which itself is integral to the use of phages within a phage therapy setting. Moreover, accurate phage genomes provide better understanding of the nucleotide and proteomic structure and how they fit into current taxonomic classification of phages. This is particularly important when performing comparative genomic analyses. We acknowledge that the production of high-quality phage genomes using this workflow requires sequencing and bioinformatic capabilities, and may be a limiting factor for some.

## AUTHOR CONTRIBUTIONS

Conceptualisation: CKAE, TLB, EMA.

Data curation: CKAE, TLV, RE, DJB, SKT, GT, HAK, RAK, LJH.

Formal analysis: CKAE, EMA.

Funding acquisition: EMA.

Investigation: CKAE, TLV, GT, HAK.

Methodology: CKAE, TLB, TLV, AT, EMA.

Software: TLV, AT.

Supervision: EMA, MAW.

Validation: CKAE, TLV, EMA.

Visualisation: CKAE.

Writing – original draft: CKAE.

Writing – review and editing: TLB, TLV, RE, DJB, AT, SKT, HAK, GT, RAK, LJH, MAW, EMA.

## ACKNOWLEDGEMENTS

We would like to thank Dr Oliver Charity for assistance with comparative genomics. We gratefully acknowledge CLIMB-BIG-DATA infrastructure (MR/T030062/1) support for high-performance computing. We would like to thank Dr Kata Farkas and Prof Davey Jones at Bangor University for their assistance in procuring wastewater samples. We would like to thank Dr James Soothill at Great Ormond Street Hospital for additional clinical *Klebsiella* strains. We would like to thank all other members of the Adriaenssens Group, including Luke Acton at Quadram Institute Bioscience for their feedback and support.

## ETHICAL STATEMENT

An ethical statement for this study was not necessary since no clinical samples were processed. However, the ethical statement for the preterm infant isolates is available from the original study (25).

## Abbreviations

BHI: brain heart infusion
CPS: capsular polysaccharide
DNAP: DNA polymerase
DTR: direct terminal repeat
HYPPA: hybrid and poly-polish phage assembly
ICTV: International Committee on Taxonomy of Viruses
LPS: lipopolysaccharide
NCBI: National Centre for Biotechnology Information
MLST: multilocus sequence typing
ONT: Oxford Nanopore Technologies
PEG: polyethylene glycol
QC: quality control
QIB: Quadram Institute Bioscience
RNAP: RNA polymerase

