## Supplemental Data for "A hybrid and poly-polish workflow for the complete and accurate assembly of phage genomes: a case study of ten przondoviruses" (from another version of this preprint: version 1, 2023-03-09)

### SUPPLEMENTAL INFORMATION

#### Supplemental tables

**Table S1.** Metadata for bacterial strain collection.

| Bacterial strains | Code or Description | Source | K locus type | MLST | Accession | Reference |
|---|---|---|---|---|---|---|
| Clinical isolates | | | | | | |
| *K. aerogenes* | CL1 | Child | - | - | - | - |
| *K. aerogenes* | CL2 | Child | - | - | - | - |
| *K. aerogenes* | CL3 | Child | - | - | - | - |
| *K. grimontii* | P038I | Infant faeces | None | None | QFVM00000000 | (25) |
| *K. grimontii* | P043G P | Infant faeces | KL169 | ST216 | QFVL00000000 | (25) |
| *K. michiganensis* | M7 21 2 #35 | Infant faeces | None | ST85 | - | - |
| *K. michiganensis* | P049A W | Infant faeces | KL102 | None | QFUG00000000 | (25) |
| *K. michiganensis* | P095L Y | Infant faeces | KL109 | ST157 | QFTT00000000 | (25) |
| *K. oxytoca* | M59 22 8 KoN | Infant faeces | KL74 | ST215 | - | - |
| *K. oxytoca* | M59 22 8 KoA | Derived from KoN | KL74 | ST215 | - | - |
| *K. oxytoca* | CL4 | Child | - | - | - | - |
| *K. oxytoca* | CL5 | Child | - | - | - | - |
| *K. pneumoniae* | CL6 | Child | - | - | - | - |
| *K. pneumoniae* | CL7 | Child | - | - | - | - |
| *K. pneumoniae* | M26 18 1 | Infant faeces | None | ST432 (CG1777) | - | - |
| *K. pneumoniae* | M26 18 2 #21 KpnN | Derived from M26 18 1 | None | ST432 (CG1777) | - | - |
| *K. pneumoniae* | M26 18 2 #21 KpnA | Derived from KpnN | None | ST432 (CG1777) | - | - |
| *K. pneumoniae* | P008E 1 | Infant faeces | KL10 | ST461 | QFUX00000000 | (25) |
| *K. quasipneumoniae* | P057K W | Infant faeces | KL11 | ST4326-2LV | QFVB00000000 | (25) |

| | | | | | | |
|---|---|---|---|---|---|---|
| *K. variicola* | CL8 | Child | - | - | - | - |
| Food isolates* | | | | | | |
| *K. pneumoniae* | 133-137-25 | Chicken meat product | KL8 | ST453 | SAMN33593347 | - |
| *K. pneumoniae* | 148-152-26 | Chicken meat product | KL21 | ST290 | SAMN33593348 | - |
| *K. pneumoniae* | 0311-342-6 | Pork meat product | KL24 | ST15 (CG15) | SAMN33593349 | - |
| *K. pneumoniae* | H-26-2393-2397-33 | Pork meat product | None | ST15 (CG15) | SAMN33593350 | - |
| *K. pneumoniae* | H-27-MC778-782col38B | Chicken meat product | KL16 | ST14 (CG15) | SAMN33593351 | - |
| Misc. isolates | | | | | | |
| *K. pneumoniae* | ST38 01 | Unknown | None | ST134 (CG1) | - | - |
| Type isolates | | | | | | |
| *K. aerogenes* | DSM 30053 | DSMZ | None | ST134 (CG1) | CP002824 | (26) |
| *K. pneumoniae* | DSM 30104 | DSMZ | KL3 | ST3 (CG3) | AJJI01000001 | (27) |
| *K. pneumoniae* | NC 13438 | NCTC | KL107 | ST258 (CG258) | UGKS01000005 | (28) |
| *K. variicola* | DSM 15968 | DSMZ | KL16 | ST2263 | CP010523 | (29) |
| Wastewater isolates | | | | | | |
| *K. huaxiensis* | 0220.RS02 | Raw sewage | None | None | - | This study |
| *K. michiganensis* | 0220.ML02 | Mixed liquor | None | None | - | This study |
| *K. michiganensis* | 0220.ML04 | Mixed liquor | None | None | - | This study |
| *K. pneumoniae* | 0220.ML03 | Mixed liquor | KL38 | ST37 (CG37) | - | This study |
| *K. pneumoniae* | 0220.RS01 | Raw Sewage | KL30 | ST1618 | - | This study |
| *K. ornithinolytica* | 0220.AF01 | Aeration feed | None | None | - | This study |
| *K. ornithinolytica* | 0220.ML01 | Mixed liquor | KL144 | None | - | This study |

*, food isolates overall BioProject accession PRJNA941224; MLST, multilocus sequence type; ST, sequence type; CG, clonal group; -, no data.

**Table S2.** HYPPA assembly details for all ten przondoviruses in this study.

| Phage lab code | Phage name | Assembler | No. contigs | No. of reads | Read length N50 | Longest read (bp) | Long-read coverage | Size pre-polishing (bp) | Size post-polishing* (bp) | Final assembly (bp) | DTR size (bp) |
|---|---|---|---|---|---|---|---|---|---|---|---|
| CE3 | Oda | Canu | 1 | 1,825 | 12,634 | 41,869 | 146x | 41,761 | 41,769 | 41,642 | 181 |
| CE8 | Toyotomi | Flye | 1 | 365,482 | 39,443 | 106,111 | 117,981x | 41,249 | 41,286 | 41,268 | 180 |
| CE9 | Mera | Flye | 3[a] | 13,252 | 12,497 | 58,219 | 8x | 41,454 | 40,619 | 41,400 | 180 |
| CE10 | Speegle | Flye | 1 | 47,260 | 12,610 | 81,712 | 23x | 41,394 | 41,395 | 41,395 | 180 |
| CE11 | Cornelius | Canu | 1 | 5,719 | 1,388 | 47,884 | 118x | 40,555 | 40,564 | 40,437 | 180 |
| CE12 | Tokugawa | Canu | 1 | 1,018 | 19,093 | 41,581 | 92x | 41,532 | 41,542 | 41,414 | 181 |
| CE15 | Saitama | Canu | 1 | 19,217 | 2,597 | 43,752 | 661x | 46,600 | 46,613 | 40,741 | 181 |
| CE20 | Emom | Canu | 1 | 2,683 | 4,862 | 41,521 | 52x | 40,903 | 40,915 | 40,788 | 183 |
| CE22 | Amrap | Flye | 2[a] | 30,181 | 717 | 40,963 | 27x | 41,328 | 41,331 | 41,209 | 182 |
| CE30 | Whistle | Canu | 1 | 9,239 | 1,614 | 46,197 | 200x | 40,855 | 40,862 | 40,735 | 181 |

*, Post-polishing was 2 x long-read and 2 x short-read polishing, as per the HYPPA workflow; [a], genome contained within the first contig

**Table S3.** Alternative long-read-only assembly details for all ten przondoviruses in this study.

| Phage lab code | Phage name | Assembler | No. contigs | Size (bp) | Size post-polishing (bp) |
|---|---|---|---|---|---|
| CE3 | Oda | Flye | 1 | 41,116 | 41,119 |
| CE8 | Toyotomi | N/A | N/A | N/A | N/A |
| CE9 | Mera | Canu | 1 | 41,527 | 40,237 |
| CE10 | Speegle | Canu | 5 | 31,413[a] | N/A |
| CE11 | Cornelius | Flye | 1 | 40,487 | N/A |
| CE12 | Tokugawa | Flye | 1 | 40,791 | 40,796 |
| CE15 | Saitama | Flye | 6 | 40,551[b] | 40,558 |
| CE20 | Emom | Flye | 1 | 41,250 | N/A |
| CE22 | Amrap | Canu | 1 | 25,100[a] | N/A |
| CE30 | Whistle | Flye | 1 | 40,843 | N/A |

N/A, not performed; [a], incomplete assembly; [b], genome contained within the first contig

**Table S4.** Short-read-only assembly details for all ten przondoviruses in this study.

| Phage lab code | Phage name | Assembler | No. contigs | No. reads | Short-read coverage | Size (bp) | Size post-curation (bp) |
|---|---|---|---|---|---|---|---|
| CE3 | Oda | Shovill | 1 | 527,773 | x3737 | 41,540 | 41,641 |
| CE8 | Toyotomi | Shovill | 46 | 396,472 | x2832 | * | N/A |
| CE9 | Mera | Shovill | 1 | 102,411 | x737 | 41,315 | N/A |
| CE10 | Speegle | SPAdes | 56 | 55,213 | x401 | 41,270 | N/A |
| CE11 | Cornelius | Shovill | 1 | 353,905 | x2608 | 40,336 | N/A |
| CE12 | Tokugawa | Shovill | 1 | 414,141 | x2935 | 41,312 | 41,413 |
| CE15 | Saitama | Shovill | 4 | 1,181,498 | x256 | 40,633 | N/A |
| CE20 | Emom | Shovill | 1 | 1,027,070 | x5627 | 40,692 | N/A |
| CE22 | Amrap | Shovill | 1 | 902,433 | x1846 | 41,106 | N/A |
| CE30 | Whistle | Shovill | 1 | 1,914,368 | x13,511 | 40,641 | N/A |

N/A, not performed; *, no single contig contained the entire genome for CE8: assembly was performed with subsampled reads in multiple iterations, but this did not improve the assembly quality.

**Table S5.** Hybrid assembly details for all ten przondoviruses in this study.

| Phage lab code | Phage name | Assembler | No. contigs | Size (bp) |
|---|---|---|---|---|
| CE3 | Oda | N/A | N/A | N/A |
| CE8 | Toyotomi | SPAdes<br>Unicycler | 64<br>6 | 40,626[a]<br>40,563[a] |
| CE9 | Mera | SPAdes | 124 | 41,329[b] |
| CE10 | Speegle | SPAdes | 74 | 41,270[c] |
| CE11 | Cornelius | N/A | N/A | N/A |
| CE12 | Tokugawa | N/A | N/A | N/A |
| CE15 | Saitama | N/A | N/A | N/A |
| CE20 | Emom | N/A | N/A | N/A |
| CE22 | Amrap | N/A | N/A | N/A |
| CE30 | Whistle | N/A | N/A | N/A |

N/A, not performed; [a], genome contained within the first two and largest contigs; [b], genome contained within the first seven and largest contigs; [c], genome contained with the first contig.

**Table S6.** Metadata for a selection of publicly-available *Autographiviridae* phages downloaded from NCBI for comparative genomics.

| Phage name | Accession | Genome size (bp) | DTR (bp)[a] | Reference |
|---|---|---|---|---|
| *Studiervirinae*, *Przondovirus* | | | | |
| *Escherichia* phage K30 | HM480846 | 40,940 | Yes (393) | (67) |
| *Klebsiella* phage 2044-307w | MF285615 | 40,048 | Yes (181) | - |
| *Klebsiella* phage Henu1 | MK203841 | 40,352 | No | (68) |
| *Klebsiella* phage K5 | KR149291 | 41,698 | Yes (392) | - |
| *Klebsiella* phage K11 | EU734173 | 41,181 | Yes (180) | (69) |
| *Klebsiella* phage KP32 | GQ413937 | 41,119 | No | (20) |
| *Klebsiella* phage KPN3[b] | MN101227 | 38,503 | No | - |
| *Klebsiella* phage Kund-ULIP47 | MK380015 | 41,397 | Yes (180) | (70) |
| *Klebsiella* phage Kp_Pokalde_002 | MT425185 | 41,816 | Yes (180) | |
| *Klebsiella* phage SH-KP152226 | MK903728 | 41,420 | Yes (180) | (71) |
| *Klebsiella* phage vB_KpnP_PRA33 | KY652723 | 40,605 | No | (72) |
| *Klebsiella* phage vB_KpnP_BIS33 | KY652725 | 41,697 | No | (72) |
| *Klebsiella* phage vB_KpnP_KpV767 | KX712070 | 40,395 | Yes (180) | - |
| *Klebsiella* phage KMI1[b] | MN052874 | 37,414 | No | - |
| *Klebsiella* phage SH-Kp 152410 | MG835568 | 40,945 | No | - |
| *Klebsiella* phage K5-2 | KY389315 | 41,116 | Yes (177) | (66) |
| *Klebsiella* phage K5-4 | KY389316 | 40,163 | Yes (179) | (66) |
| *Klebsiella* phage vB_Kpn-VAC71 | MZ571832 | 40,388 | No | - |
| *Klebsiella* phage vB_KpnP_IME205 | KU183006 | 41,310 | No | (73) |
| *Klebsiella* phage IME264 | OL799328 | 40,671 | No | - |
| *Studiervirinae*, *Teetrevirus* | | | | |
| *Salmonella* phage phiSG-JL2 | EU547803 | 38,815 | Yes (230) | (74) |
| *Citrobacter* phage SH1 | KU687347 | 39,434 | Yes (231) | (75) |
| *Klebsiella* phage vB_KpnP-VAC1 | MZ428229 | 39,371 | No | (76) |
| *Studiervirinae*, *Teseptimavirus* | | | | |
| *Escherichia* phage T7 | V01146 | 39,937 | Yes (160) | (77) |
| Enterobacteria phage 13a | EU734174 | 38,841 | Yes (170) | - |
| *Studiervirinae*, *Apidecimavirus* | | | | |
| *Yersinia* phage vB_YenP_AP10 | KT852574 | 39,235 | Yes (193) | - |
| *Studiervirinae*, *Berlinvirus* | | | | |
| *Yersinia* phage Berlin | AM183667 | 38,564 | Yes (226) | - |
| *Slopekvirinae*, *Drulisvirus* | | | | |
| *Klebsiella* phage F19 | KF765493 | 43,766 | Yes (119)[c] | - |
| *Molineuxvirinae*, *Zindervirus* | | | | |
| *Salmonella* phage SP6 | AY370673 | 43,769 | Yes (174) | (78) |

[a], Yes, complete genomes were those with the DTRs present that may or may not be annotated; No, genomes without DTRs present and therefore deemed incomplete. [b], Klebsiella phages KPN3 and KMNI1 were deemed incomplete due to potential errors in multiple genes. [c], DTRs were annotated but only the first 94 bp of the 119 bp annotation are homologous. -, no data.

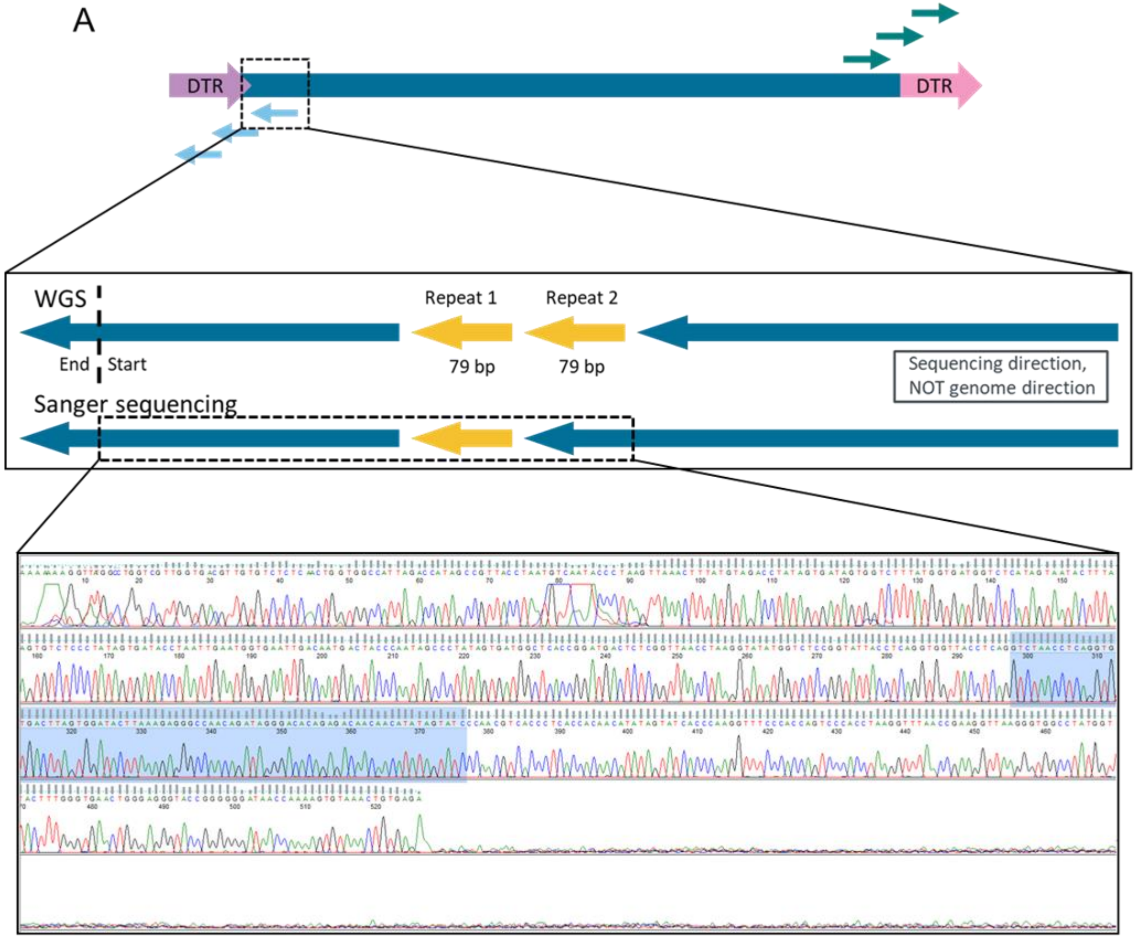

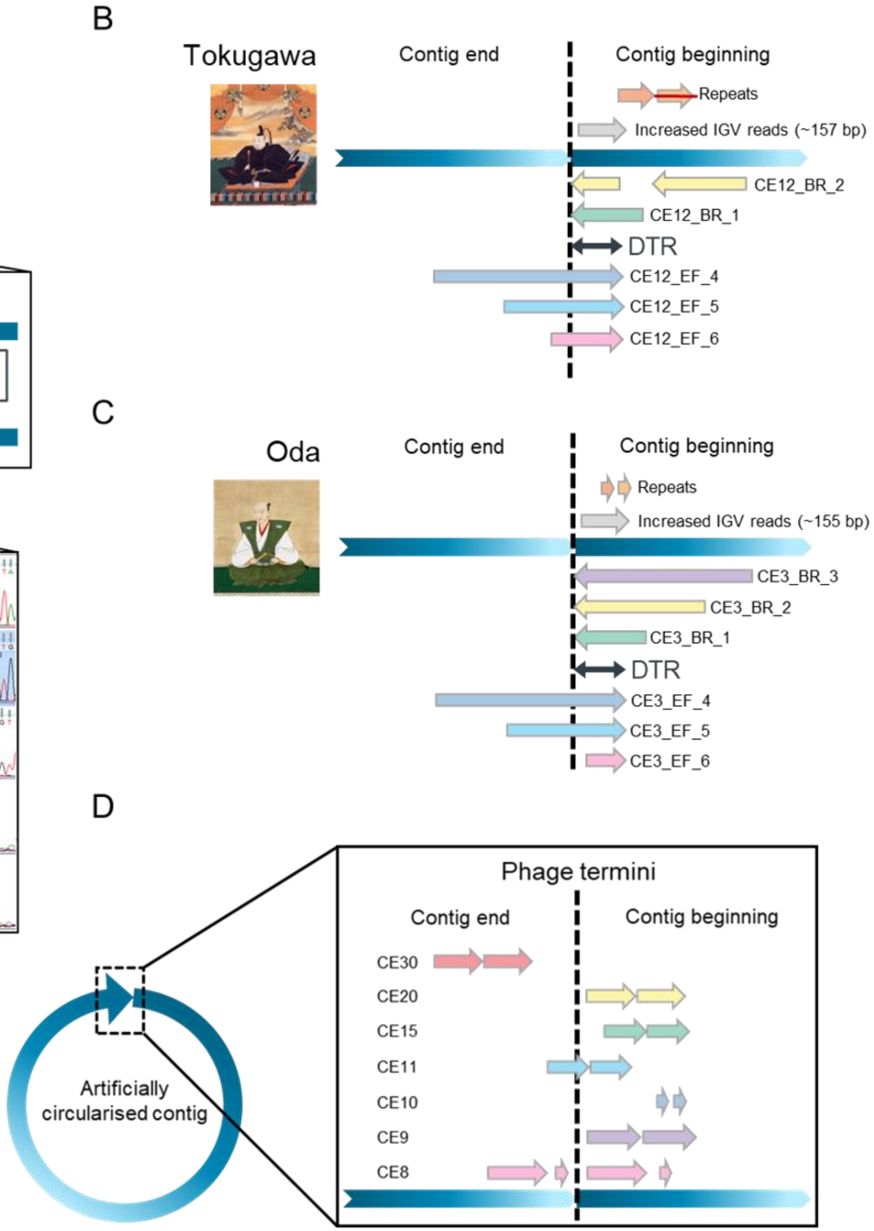


**Fig. S2. Schematic of primer walking and Sanger sequencing validation of the HYPPA workflow. (A)** Three primers were designed to “walk” the phage termini (top panel). Results for *Klebsiella* phage Tokugawa showed that there were two 79 bp repeats in the short read - only assembly data (middle panel), where only one was present in the Sanger sequencing data (middle and bottom panel), as is highlighted in the chromatograph, suggesting an assembly error. **(B)** Schematic of the primer walking validation for *Klebsiella* phage Tokugawa showing where the 79 bp repeats were, and that one was introduced in error. The two reverse and three forward primers marked the area where the DTR began and ended, and corresponded to the area of increased IGV reads. **(C)** Schematic of the primer walking validation for *Klebsiella* phage Oda using three primers for each terminus. The assembly had small 13 bp repeats that were confirmed as genuine following Sanger sequencing. **(D)** Schematic of the phage termini with all the larger repeats introduced in error during short-read-only assembly. The small repeats found in CE8 and CE10 were genuine. small repeats of 13 bp.

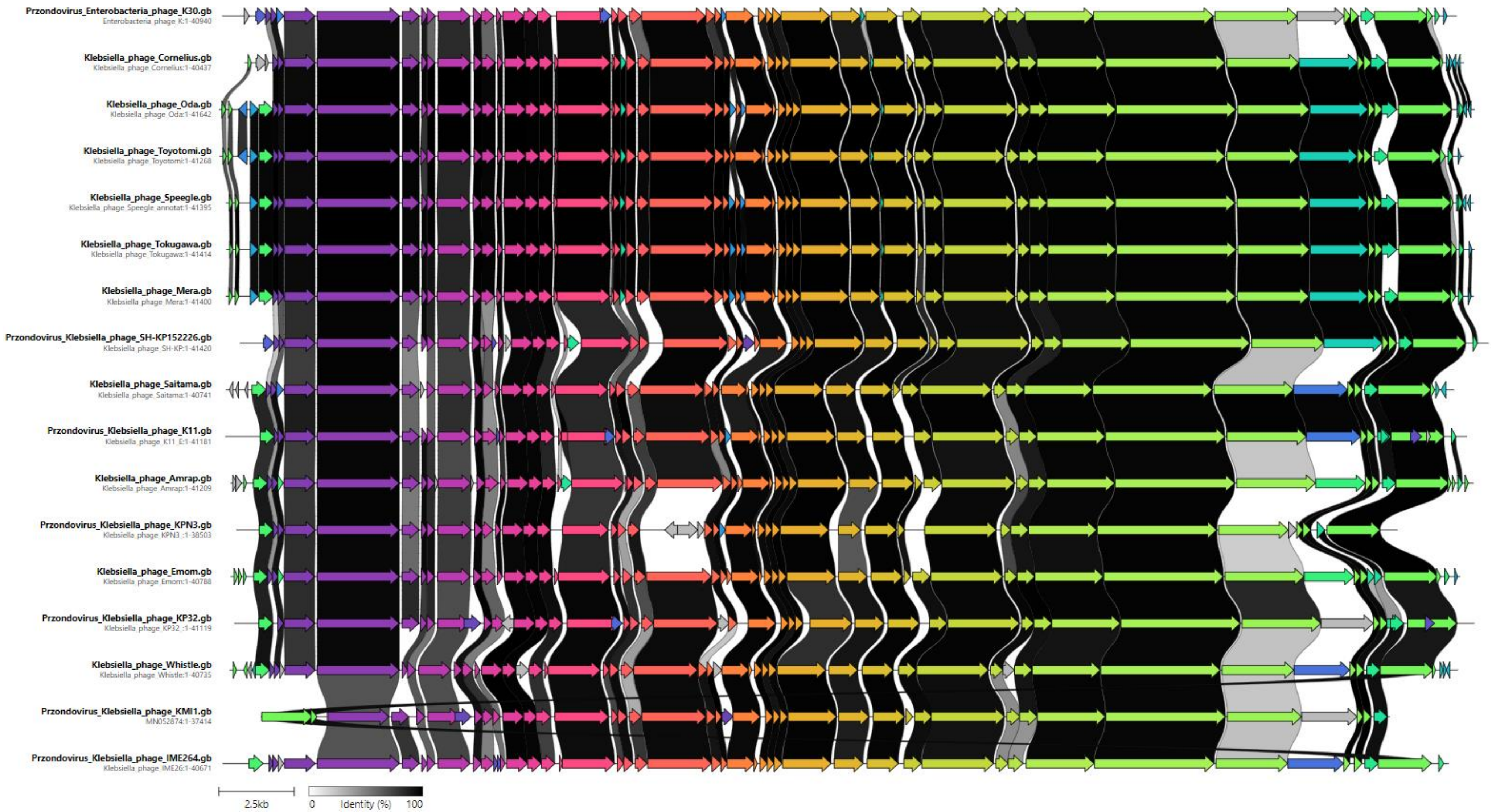


**Fig. S2. Genome map and gene clustering for przondoviruses in the collection and a selection of related phages including relatives with potential errors.** Arrows represent coding sequences and pairwise comparisons of gene similarities are indicated by percentage identity given as links in greyscale, with darker shading representing areas of higher similarity. Genes without any sequence similarity are indicated without links. DTRs are not annotated. Potential assembly/sequencing errors in reference phages: i) several missing genes including S-adenosyl-L-methionine hydrolase in phage SH-KP152226; ii) incomplete annotation of the DNA polymerase and missing tail tubular protein A in phage KPN3; iii) incorrect start site of phage KMI1, along with incomplete DNA polymerase and several missing genes, including S-adenosyl-L-methionine hydrolase and serine/threonine kinase.
